# Coupled phylogenetic and functional enrichment in the tomato rhizosphere microbiome

**DOI:** 10.1101/2024.05.22.595324

**Authors:** Silvia Talavera-Marcos, Ramón Gallego, Rubén Chaboy, Alberto Rastrojo, Daniel Aguirre de Cárcer

## Abstract

Plant-microbe interactions occur mainly in the rhizosphere, a hot spot of microbial activity and diversity. Given that the outcome of such interactions can significantly impact plant productivity, we require a better understanding of the rhizosphere microbiome if knowledge-based microbiome modification strategies are to be successfully deployed in the future. Here, we aimed to gain a better understanding of the assembly process of the tomato rhizosphere microbiome and its potential composition-function relationships. Among other things, we studied community assembly through the lens of a conceptual framework for the phylogenetically constrained assembly of microbial communities, while assessing community function based on the predicted minimal metagenome of the microbial ecosystem. We observed a systematic enrichment in terms of phylogeny and predicted functional content in the rhizosphere and were able to delimit phylogenetic signal in the ecosystem with 12 functionally coherent phylogenetic groups present in all samples which together accounted for a large fraction of the total community. Our analyses indicated that these groups included a significantly larger content of the ecosystem’s minimal metagenome than expected by chance. Thus, our study suggests that community assembly followed coupled phylo-functional selection independent of host genetics, and we expect the same phenomenon to occur in other rhizosphere microbiomes. This knowledge provides a thrust in our understanding of how community composition-phylogeny-function relationships drive the assembly process of the rhizosphere microbiome and should help guide the design of synthetic rhizosphere microbiomes for both research and commercial purposes.

---

The interface between plant root and soil, the rhizosphere, is a hot spot of microbial activity and diversity, and is considered to be among the most complex microbial ecosystems on the planet (Raaijmakers et al. 2009). In this millimeter-scale environment, plants actively select and support a wide array of microbes through specific root exudates and cell debris (Bais et al. 2004; Philippot et al. 2013). In return, the microbial inhabitants of the rhizosphere can promote plant fitness by enhancing nutrient bioavailability or the production of phytohormones (Backer et al. 2018), increasing tolerance to abiotic stress factors (Yang et al. 2009; Berendsen et al. 2012), or suppressing deleterious microorganisms through antagonistic interactions or competitive exclusion (Mendes et al. 2013).

Mimicking the strong interplay between the human gut microbiome and its host (Visconti et al. 2019), plant-microbe interactions occur mainly in the rhizosphere (Berendsen et al. 2012), where intensive plant-microbe and microbe-microbe interactions take place (Philippot et al. 2013). Given that the outcome of such interactions can significantly impact plant productivity, and the urgent need to transition to a more sustainable agriculture where microbiome-derived products and services are expected to play a significant role, there is a vital need to increase our understanding of the rhizosphere microbiome. Thus, in addition to an increase in our mechanistic comprehension of microbe-microbe and plant-microbe interactions, we require a better understanding of the assembly process of the rhizosphere microbiome and its community composition-function relationships if knowledge-based microbiome modification strategies are to be successfully deployed in the near future.

Tomato (*Solanum lycopersicum*) is an important crop plant around the globe and is also employed as a research model (Bai et al. 2018). As such, the study of its associated microbiome has increased in the last few years, providing us with valuable information on its community composition in different plant compartments (Ottesen et al. 2013; Dong et al. 2019), knowledge of the protein diversity present in its rhizosphere (Barajas et al. 2020), and an understanding of the interplay between host genetics and microbiome assembly (Oyserman et al. 2022). The aim of this study is to gain a better understanding of the assembly process of the tomato rhizosphere microbiome and its potential composition-function relationships. To do so, we studied community assembly through the lens of a conceptual framework for the phylogenetically constrained assembly of microbial communities (Aguirre de Cárcer 2019). The framework focuses firstly on the tendency of most microbial communities to show phylogenetic clustering (i.e. when bacteria tend to co-occur with phylogenetically related populations more often than expected by chance), and the fact that such phylogenetic signal in microbial communities can be studied in terms of phylogenetic core groups (PCGs); specific nodes in the bacterial phylogeny present in all instances of a given ecosystem type. Secondly, it also draws from the fact that the bacterial 16S rRNA gene-based phylogeny presents significant functional coherence (i.e. within-node gene content similarity is significantly higher than expected by chance) (Parras-Moltó and Aguirre de Cárcer 2021) The framework proposes that the most reasonable explanation for the existence of a PCG in a given environment is that its intra-PCG populations present a phylogenetically conserved set of traits that improve the fitness of those populations under the biotic and abiotic factors in that environment. Thus, the existence of a PCG could be understood as likely related to a specific niche whose occupancy requires a specific phylogenetically conserved set of traits.

In a previous study, we showed that the existence of PCGs was a predominant feature of the wide array of microbial ecosystems studied (Talavera-Marcos et al. 2023), which included human, animal, plant, and environmental microbiomes. Furthermore, we analyzed of an *in vitro* experimental dataset using genome-scale metabolic models and population dynamics modelling and obtained ecological insights on metabolic niche structure and population dynamics comparable to those gained after canonical experimentation (Estrela et al. 2022), thus demonstrating the potential of leveraging phylogenetic signal to help unravel microbiome function and assembly rules.

In the present work, we applied the framework to the study of the tomato rhizosphere. The experimental design featured high replication, employing up to eight replicates for each of the eight different starting communities, so that the phylogenetic signal of the tomato rhizosphere could be adequately delimited. The plant host originated from a single, genetically homogenous variety to avoid the expected noise that would derive from the use of different plant genotypes (Oyserman et al. 2022). The in planta experimental scenario precludes the use of the modelling approaches employed in our previous study, since the available carbon and energy sources in the ecosystem are so far not sufficiently characterized. Instead, we evaluate community function using a different perspective based on the analysis of the predicted metagenomic content of the microbial ecosystem.

Tomato seeds (*Salinas F1*, CapGen Seeds (Spain)) were surface sterilized by immersion in 70% ethanol for 2 min and 10% bleach for another 2 min, washed four consecutive times with sterile distilled water, and finally incubated for 1 h on sterile distilled water. The resulting seeds were pre-germinated on sterile 1.5% agar plates for two days at 28ºC in the dark before they were sown on 50 ml polypropylene tubes containing a mixture of 6 g of washed sterile perlite and 3 g of soil and irrigated with 5 ml FP plant mineral medium (Fahraeus 1957). The 8 different soils employed arose from different locations in Spain (A; Recreational orchard from Guadalajara province. B; Recreational orchard from Madrid province. C; Forest soil from Madrid province. D; Recreational Orchard from Pontevedra province. E; Recreational Orchard from Cantabria province. F; Ruderal soil from Alicante province. G; Dryland cultivation soil from Badajoz province. H; Ruderal soil from Madrid province). Topsoil samples from each location were sieved through a clean 1 mm mesh to remove large debris before planting.

Eight plants per soil type were grown for three weeks under controlled conditions (24 ºC and 16 h light followed by 18 ºC and 12 h darkness) before sampling. Using clean forceps, root systems were vigorously shaken to remove loosely attached soil particles, and then introduced into 2 ml polypropylene tubes containing 1 ml ice-cold sterile PBS buffer. The tubes were vortexed for 5 min to resuspend the rhizosphere community. Three *ca*. 300 g replicates from each soil type and 300 µl of each rhizosphere community were subsequently processed for microbial community analysis.

DNA was extracted using the *MagBind Environmental DNA 96 Kit* (Omega BioTek) according to the manufacturer’s instructions. The quality of extracted DNA was checked through agarose gel electrophoresis and its concentration measured using *Picogreen* (Invitrogen). The 16S rRNA bacterial phylogenetic marker gene was amplified from each resulting DNA sample using primers 347F and 803R targeting the V3-V4 hypervariable region and including Illumina sequencing adapters. Amplification was performed using 4 µl of template DNA in a 44 µl reaction mixture containing 0.06 µM of each primer, 0.5 µM dNTPs and 1 U of *Q5 HighFidelity* DNA Polymerase (New England Biolabs). The thermocycler conditions consisted of 95 °C for 30 s, followed by 20 cycles of 95 °C for 10 s, 55 °C for 30 s, and 72 °C for 30 s. PCR products were then treated with 2 µl of a 1X *Q5 HighFidelity* DNA Polymerase buffer and 5 U/µl *ExoI* exonuclease (Thermo Scientific) solution for 20 min at 37ºC and 15min at 80ºC to remove the initial primers. The resulting products were then subjected to 10 more PCR cycles as above but adding 3 µl of 10 mM primer stocks bearing (5’-3’) the required i5 and i7 Illumina adapters, 10 nt barcodes, and Illumina sequencing primers. The amplicon libraries produced were checked through agarose gel electrophoresis and their concentration measured using *Picogreen* (Invitrogen). Equimolar amounts from each library were pooled, ran on an agarose gel, the appropriate band excised and purified using the *QIAquick Gel Extraction Kit* (Qiagen). The final product was sequenced on an Illumina MiSeq NGS platform using a 600 cycle v3 reagent kit following the manufacturer’s instructions.

Initial sequence processing with *R* package *DADA2* (Callahan et al. 2016) included its standard pipeline for error modelling, paired-end sequence merging, chimera removal, taxonomic assignments using SILVA’s training set (v123) (Quast et al. 2013), and the elimination of residual eukaryotic sequences including *Mitochondria* and *Chloroplast*-affiliated sequences. Then, sequences arising from the same soil sample were pooled before all samples were subsampled to a common depth. Subsampling depth was chosen based on the observed natural break in the distribution of the number of sequences (Schloss 2024), and samples below such depth were discarded. For analyses using phylogenetic metrics, ASVs were mapped to GreenGenes 99% representative sequences set (v13_8) using *Blast*, and its corresponding phylogenetic tree employed. Faith’s phylogenetic diversity and DPCoA were then calculated using *R* packages *biomeUtils* and *phyloseq* (McMurdie and Holmes 2013), respectively. This last package was also used for Non-Metric Multi-Dimensional analysis (NMDS) using Bray-Curtis dissimilarities, while Principal Component Analysis with respect to Instrumental Variables (PCAIV) and its associated permutation test were carried out with *R* package *ade4* (Dray and Dufour 2007). Functional metagenomic predictions and corresponding differential abundance tests were carried out using *PICRUSt2* with default parameters (Douglas et al. 2020). Phylogenetic Core Groups (PCGs) were detected using *Bacterial*C*ore*.*py* (Talavera-Marcos et al. 2023) based on GreenGenes 99% representative sequences tree (v13_8) and a minimum abundance cutoff of 0.5%. To that end, the samples originating from soil D (soil sample and the single rhizosphere sample with sufficient sequencing depth obtained) were removed, along with sequences not mapping to the reference database, and all samples were subsampled to a common depth as above. The ecological coherence of detected PCGs was assessed based on whether the detected nodes fell within the previously described limits of phylo-functional coherence along the 16S rRNA gene phylogeny (Parras-Moltó and Aguirre de Cárcer 2021). To that end, for each PCG, intra-PCG sequences were first mapped against the 16S rRNA gene set used in the original publication using *Blast*, then its last common ancestor in the accompanying phylogeny obtained using *R* package *castor*. Finally, we checked whether the last common ancestor of each PCG in that phylogeny was a descendant node from those nodes that had been described as limits of phylo-functional coherence in the original publication. All scripts and 16S rRNA gene dataset are available at https://github.com/microenvgen/tomato and the <u>European</u> Nucleotide Archive (PRJEB74653), respectively.

After the initial sample processing, 50 samples with a normalized coverage of 45000 sequences each were retained for subsequent analyses. The dataset presented a high proportion of sequences affiliated to the *Bacillaceae, Chitinophagaceae, Comamonadaceae, Cytophagaceae, Flavobacteriaceae, Oxalobacteriaceae, Pseudomonadaceae, Rhizobiaceae, Shingobacteriaceae*, and *Streptomycetaceae* families (Suppl. Fig. 1). Most commonly, soil communities presented a higher richness and phylogenetic diversity than corresponding rhizosphere samples (Suppl. Fig. 2). Such richness and diversity decrease from soil to rhizosphere is a frequently observed feature of the rhizosphere ecosystem (Dong et al. 2019; Lee et al. 2019) commonly accepted to arise from plant-derived microbiome selection (Bulgarelli et al. 2012). The exploration of microbial community profiles based on ASV compositions (Figure 1) indicated that the main driver of community structure was soil origin, and that rhizosphere communities were commonly more similar to each other than to its soil of origin. The effect of the surrounding soil on rhizosphere community structure has also been systematically reported (e.g. (Lundberg et al. 2012; Edwards et al. 2015)) as soil represents the species pool from which the rhizosphere microbiome assembles, and soil physicochemical characteristics can also influence community assembly through their impact in plan physiology (Philippot et al. 2013).

**Figure 1.**
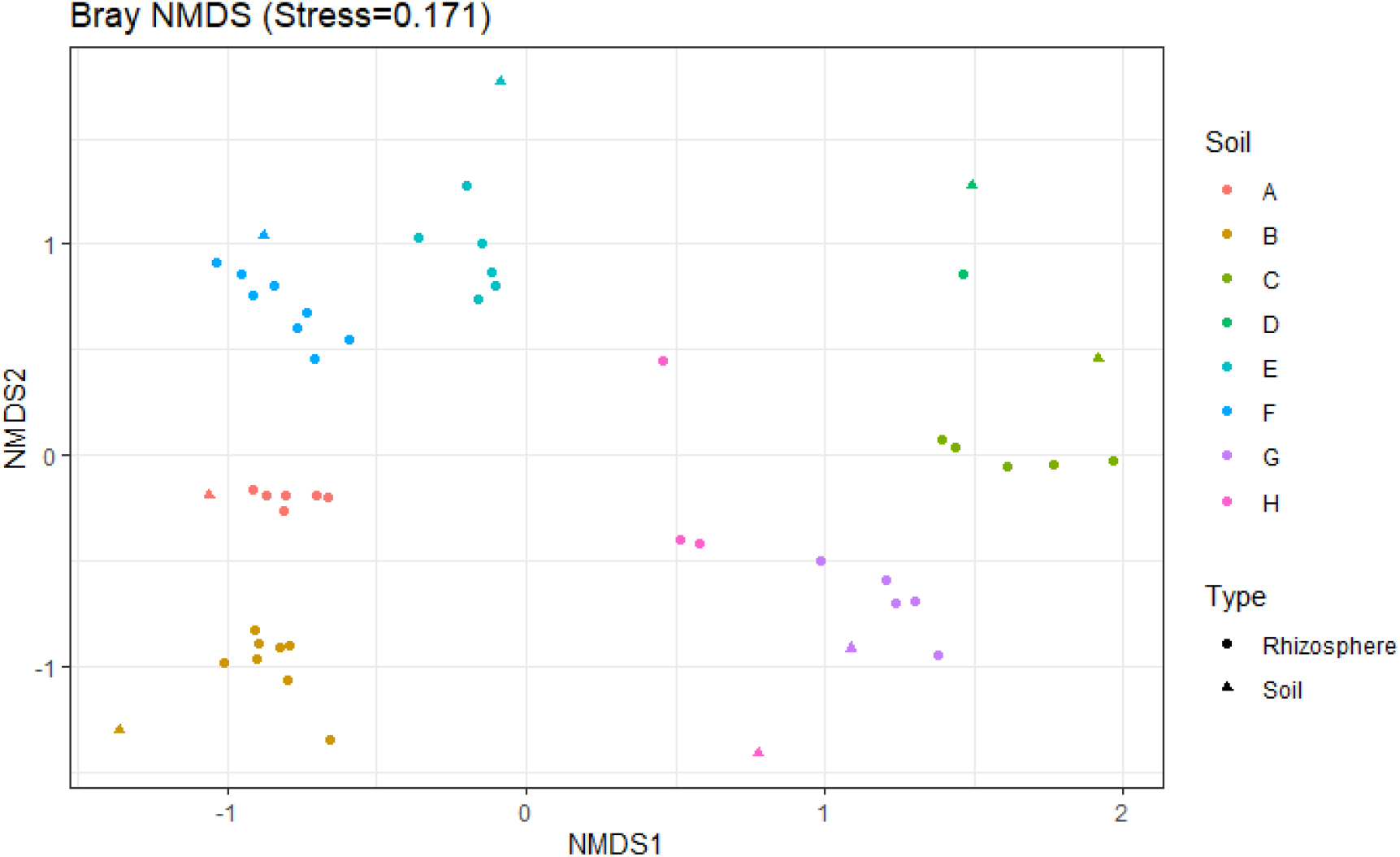
Soil of origin is the main driver of community structure. The figure depicts a non-metric multidimensional scaling plot based on Bray-Curtis distances between samples. Samples are labeled according to soil of origin by color (A-H) and sample type by shape (Rhizosphere or Soil).

However, the exploration based on phylogenetic compositions (Figure 2) showed a coordinated shift in community profiles separating rhizosphere and soil samples regardless of soil of origin. Overall, from a phylogenetic perspective, rhizosphere samples presented a higher proportion of sequences affiliated to the *Alpha, Beta*, and *Gammaproteobacteria, Flavobacteria, Sphingobacteria* and *Cytophagia*, and reduced in those affiliated to the *Bacilli, Actinobacteria* and *Acidobacteria* classes (Figure 3). To further explore this pattern, we performed a Principal Components Analysis with respect to Instrumental Variables (PCAIV) to analyze microbial community profiles based on ASV compositions after removing the soil of origin effect previously observed in Figure 1 and enforcing the exploration of differences between soil and rhizosphere samples. The results (Suppl. Fig. 3) indicated that not all ASVs affiliated to the mentioned taxonomic classes followed the observed general enrichment pattern, since, for instance, some *Acidobacteria* and *Bacilli*-affiliated ASVs were enriched in rhizosphere samples and some *Sphingobacteria* and *Alphaproteobacteria*-affiliated sequences were enriched in soil samples.

**Figure 2.**
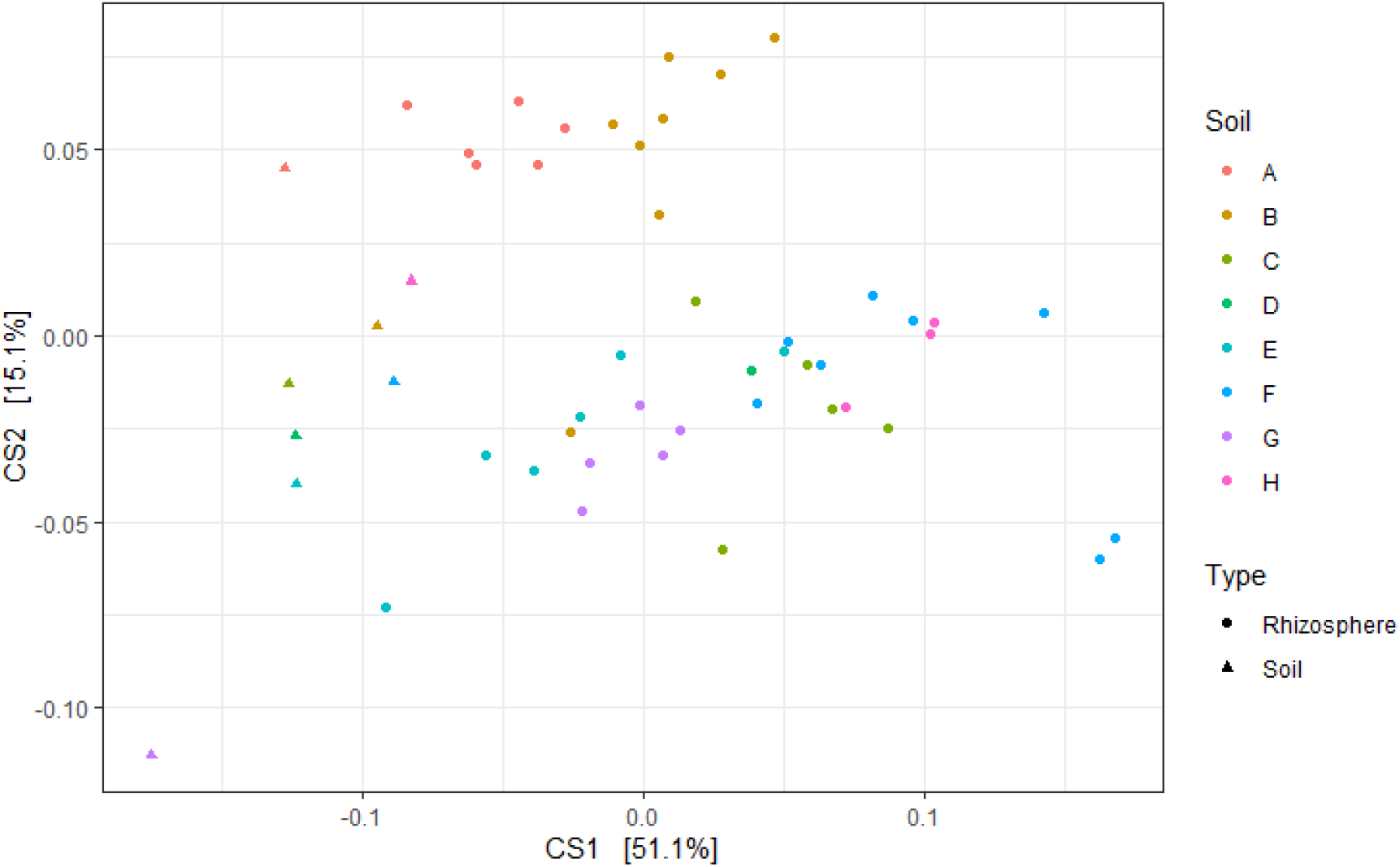
Sample type is the main driver of phylogenetic community composition. The figure depicts the first two dimensions of a double principal coordinate analysis. Samples are labeled according to soil of origin by color (A-H) and sample type by shape (Rhizosphere or Soil).

**Figure 3.**
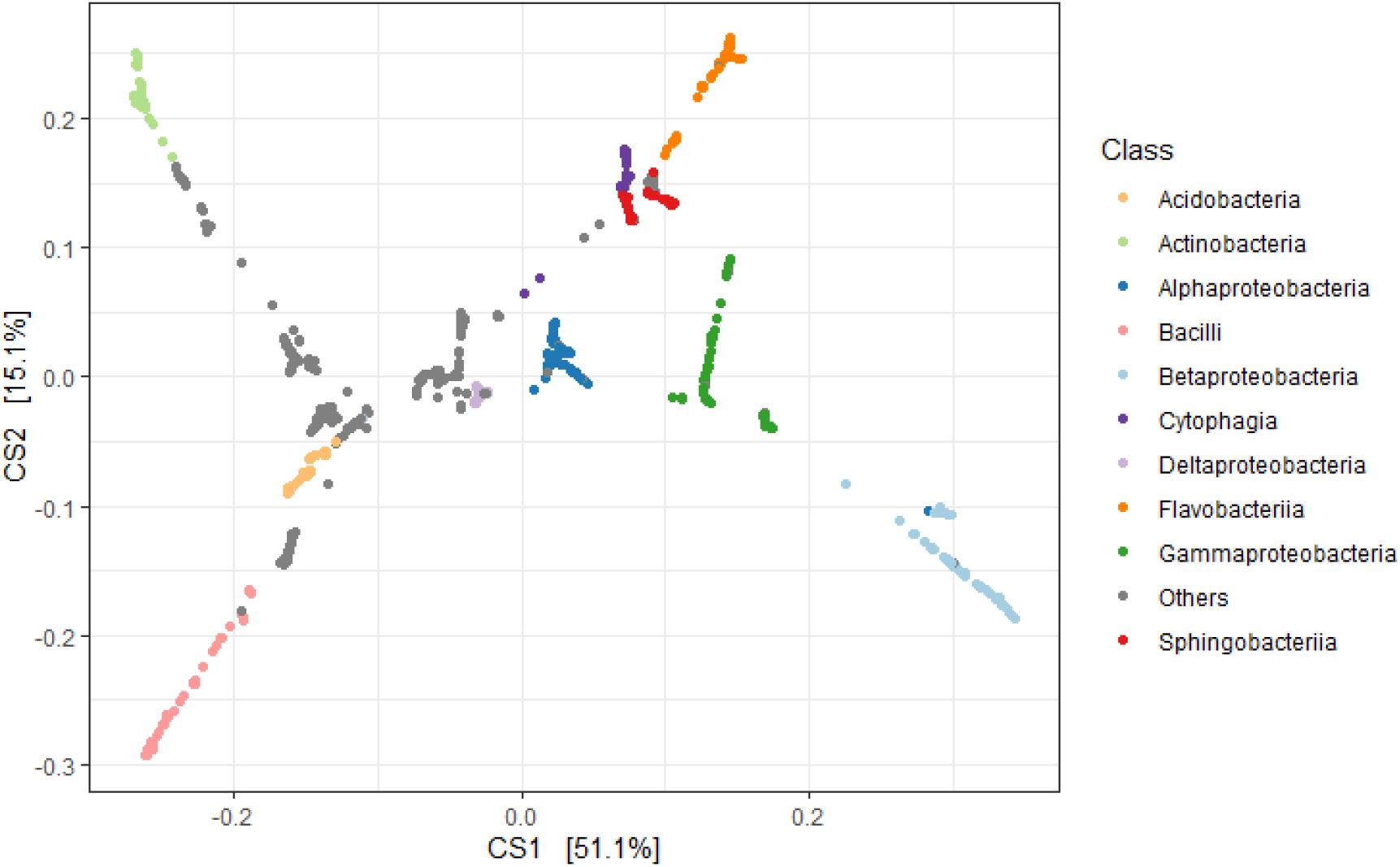
Drivers of rhizosphere-associated shift in phylogenetic community composition. The figure depicts the first two dimensions of the same double principal coordinate analysis depicted in Figure 2. Here, OTUs are labeled according to Class-level taxonomic affiliations by color (only the ten top Classes are shown).

Next, we asked whether the observed rhizosphere enrichment patterns were related to a particular set of ASVs, to the overall phylogeny as previously observed, or could also relate to overall community function. To that end, we generated functional metagenomic predictions from our 16S rRNA marker gene dataset and analyzed the resulting predictions to explore general trends in community function as well as the possible existence of significantly enriched metabolic pathways. The results also pinpointed functional enrichment for five of the eight soils of origin (Figure 4; Suppl. Fig. 4), and the overrepresentation of several metabolic pathways in rhizosphere samples, mainly related to the degradation various sugars (Glucose, Arabinose, Sucrose), amino acids (Tyrosine, Arginine, Histidine), and aromatic compounds (Benzoyl-CoA, Vanillin, Syringate, Protocaechuate, Catechol, Gallate) (Suppl. Fig. 5), common components of root exudates (McLaughlin et al. 2023).

**Figure 4.**
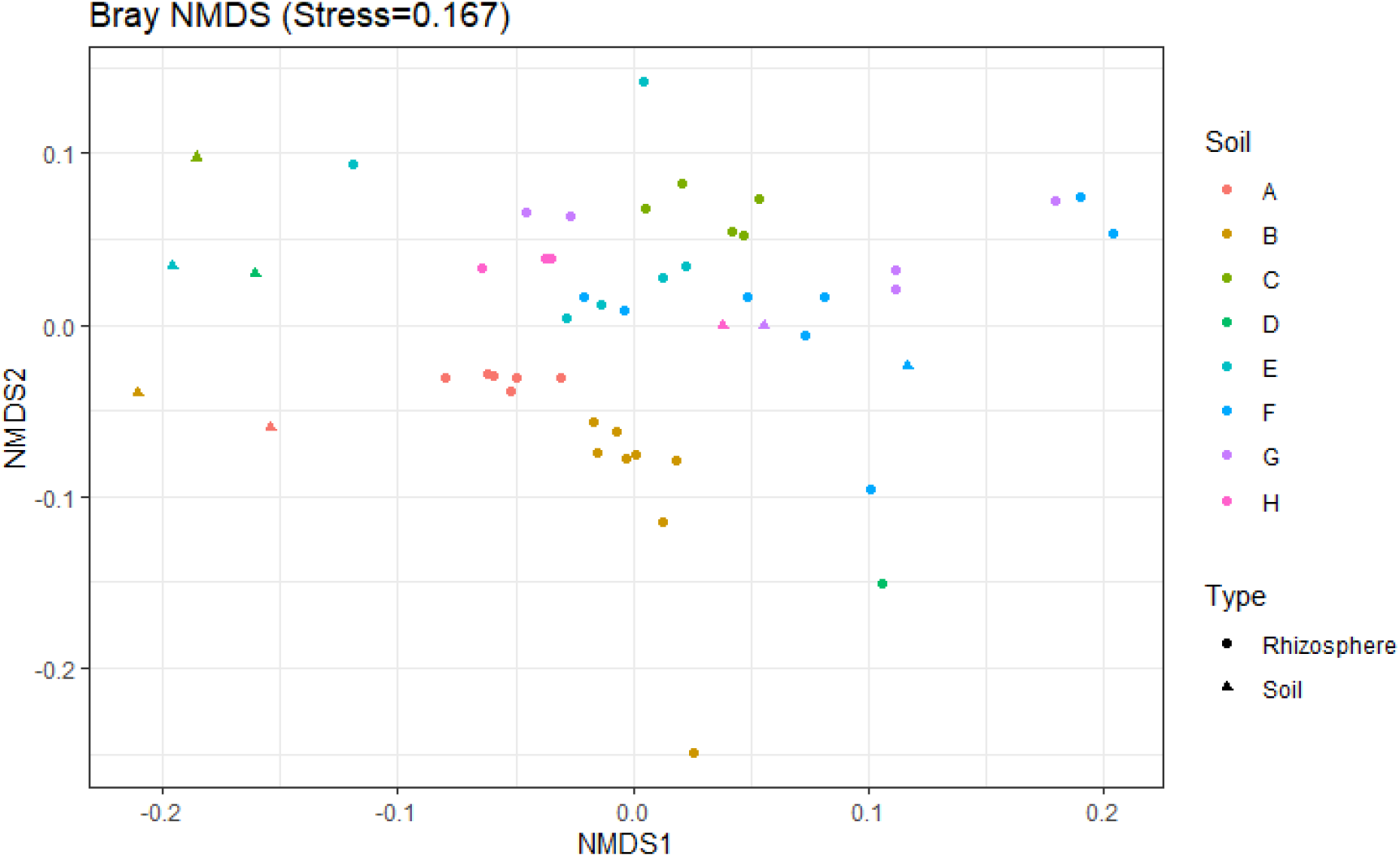
Exploration of functional community profiles. The figure depicts a non-metric multidimensional scaling based on Bray-Curtis distances between predicted EC compositions among samples. Samples are labeled according to soil of origin by color (A-H) and sample type by shape (Rhizosphere or Soil).

To further explore the possible rationale for the phylogenetic enrichment observed in the tomato rhizosphere community, we explored the existence of Phylogenetic Core Groups (PCGs) in our dataset. PCGs are discrete portions of the bacterial phylogeny present in all instances of a given microbial ecosystem type (Aguirre de Cárcer 2018), and its analysis has been proposed as a route to better understand community assembly and function (Aguirre de Cárcer 2019; Talavera-Marcos et al. 2023). According to the PCG framework, their presence would be related to the existence of specific niches in the microbial ecosystem whose occupancy requires a specific phylogenetically conserved set of traits. For the current rhizosphere dataset, we detected 20 PCGs with a minimum relative abundance of 0.5% across all our samples. Of these, 8 were detected outside of the previously-described limits of phylo-functional coherence along the 16S rRNA gene phylogeny (Parras-Moltó and Aguirre de Cárcer 2021) and thus were discarded from subsequent analyses. The resulting groups (Table 1) were affiliated to the *Alpha, Beta*, and *Gammaproteobacteria, Actinobacteria*, and *Bacilli* classes, and globally accounted for an average of *ca*. 60% of sequences in each sample.

**Table 1.** PCGs in the experimental dataset. The table provides information on the nodes detected as PCGs (PCG), the number of leaves in the overall reference tree for each PCG node (Leaves in node), the total number of leaves detected in the experiment (Leaves in experiment), the average relative abundance of each PCG along the replicates and its min-max range (Relative abundance), and the node’s consensus taxonomic affiliation and the number of leaves in the reference tree with such taxonomic affiliation (Taxonomy/Leaves in taxon).

| PCG | Leaves<br>in<br>Node | Leaves in<br>experiment | Relative<br>abundance (%) | Taxonomy/Leaves in taxon |
| --- | --- | --- | --- | --- |
| Node15327 | 2084 | 57 | 8.4 (1.1-36.7) | Comamonadaceae (F)/3036 |
| Node19398 | 6017 | 96 | 7.0 (1.0-18.8) | Betaproteobacteria (C)/10565 |
| Node21166 | 1474 | 51 | 4.6 (0.7-14.6) | Oxalobacteraceae (F)/1443 |
| Node28120 | 1620 | 29 | 5.2 (0.5-19.0) | Pseudomonas (G)/1702 |
| Node45985 | 5257 | 144 | 1.85 (0.7-4.4) | Alphaproteobacteria (F)/16849 |
| Node52227 | 1643 | 60 | 3.4 (0.8-5.4) | Sphingomonadaceae (F)/2220 |
| Node53994 | 8815 | 306 | 7.2 (1.0-17.8) | Alphaproteobacteria (F)/16849 |
| Node55420 | 1709 | 67 | 3.1 (0.76-8.7) | Rhizobiales (O)/4680 |
| Node55904 | 415 | 22 | 3.7 (0.6-9.2) | Rhizobiaceae (F)/608 |
| Node119365 | 7533 | 154 | 3.6 (0.5-10.5) | Actinomycetales (O)/11801 |
| Node126898 | 2291 | 155 | 3.5 (0.8-9.2) | Actinomycetales (O)/11801 |
| Node147305 | 5696 | 79 | 7.8 (0.5-34.3) | Bacillales (O)/8896 |

To explore the potential functional role of the PCGs in the ecosystem, we first delineated the tomato rhizosphere minimal metagenome based on metagenomic predictions. The term minimal metagenome was originally defined as the set of microbial genes necessary for the homeostasis of the whole human gut ecosystem and expected to be present in all healthy humans (Qin et al. 2010). Following the same rationale, we define here the minimal metagenome of the tomato rhizosphere as the set of functions present in all rhizosphere samples, which would be thus necessary for appropriate community function. In this study, from a total of 7604 KOs (KEGG Orthologs) predicted as present in the rhizosphere-only dataset, 5186 were present in all rhizosphere samples, thus forming the tomato rhizosphere predicted minimal metagenome (Suppl. Fig. 6).

Once furnished with a reasonable proxy for overall community function in the form of the minimal metagenome, we explored the degree to which the 12 identified functionally coherent PCGs recapitulated community function. To do so, for each of the 40 rhizosphere samples, we generated artificial communities of 12 OTUs either i) chosen at random among the OTUs detected in the sample, ii) chosen at random among the OTUs detected in the sample not belonging to any functionally coherent PCG, or iii) chosen at random from within each PCG and including only one member per PCG of those OTUs present in the sample. Following this approach, we produced 100 artificial communities per approach (i, ii and iii) and sample, thus adding to a total of 12000 simulated communities. The metagenomic predictions for these communities indicated that the PCG combinations (iii) recovered a significantly higher proportion of the tomato rhizosphere minimal metagenome compared to their corresponding randomly chosen communities (i, ii) (78.0%, 66.8% and 57.8% for (iii), (i) and (ii), respectively; t-test p-values <2.2e-16). Going one step further, we next asked which functions (KOs) in the predicted minimal metagenome were contributed exclusively by a single PCG. The results (Suppl. Table 1) indicated many such functions pertaining to a wide range of metabolic pathways. The exclusive functions included items related to important ecosystem services such as the processing of nitrogen and sulfur or methane metabolism, as well as those implicated in the processing of root-derived compounds. In many instances, different PCGs presented exclusive functions within shared metabolic pathways, thus supporting the idea that many community functions are carried out by the coordinated action of different functional groups.

In conclusion, our results indicate a systematic and explicit directional enrichment in terms of phylogeny and predicted functional content in the tomato rhizosphere. We were able to delimit phylogenetic signal in the ecosystem in terms of 12 functionally coherent PCGs, which together accounted for a large fraction of the total community. While these PCGs were affiliated to taxa commonly described as enriched in the rhizosphere (*Alpha* and *Betaproteobacteria, Bacillales, Actinomycetales*, and *Pseudomonas*), our PCG approach was able to further delimit phylogenetic enrichment to five, three and two subgroups within the *Alphaproteobacteria, Betaproteobacteria* and *Actinomycetales* (respectively), and delimit the *Bacillales*-related phylogenetic signal to an internal node that excluded 36% of its reference sequences. Furthermore, our simulations indicated that PCGs included a significantly larger content of the ecosystems predicted minimal metagenome than expected by chance. Thus, complementing Oyserman et al in-depth study on the host genetic basis of rhizosphere microbiome assembly in tomato (Oyserman et al. 2022), our study suggests that community assembly in this environment follows coupled phylo-functional selection independent of host genetics, and we expect the same phenomenon to occur in other rhizosphere microbiomes showing strong phylogenetic signal. This knowledge provides a thrust in our understanding of how community composition-phylogeny-function relationships drive the assembly process of the rhizosphere microbiome and should help guide the design of synthetic rhizosphere microbiomes for both research and commercial purposes, as well as related bacteria isolation and characterization efforts.

## Notes

### Competing Interest Statement

The authors have declared no competing interest.

