## Supplementary material 1 for "Coupled phylogenetic and functional enrichment in the tomato rhizosphere microbiome"

**Supplementary figures and table to “Coupled phylogenetic and functional enrichment in the tomato rhizosphere microbiome”**

Silvia Talavera-Marcos^a^, Ramón Gallego^a^, Alberto Rastrojo^a^, Daniel Aguirre de Cárcer^a*^

^a^Microbial and Environmental Genomics Group, Departamento de Biología, Universidad Autónoma de Madrid, Madrid, Spain.

**FIGURES**

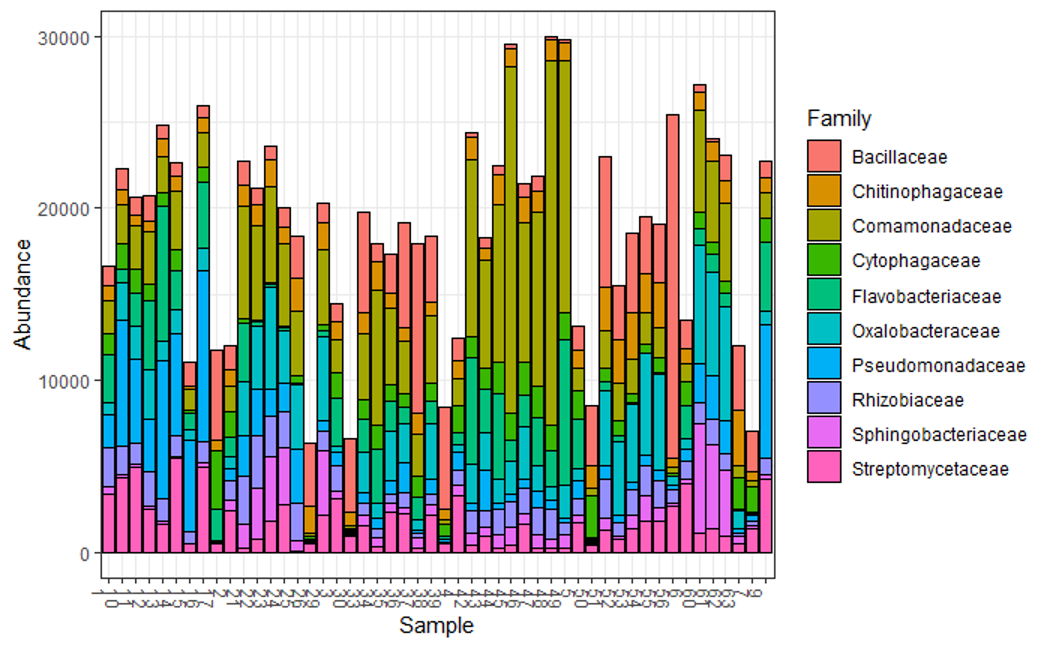

**Supplementary figure 1.** Stacked barplot representing the abundance of the top 10 most abundant taxonomic families in the dataset.

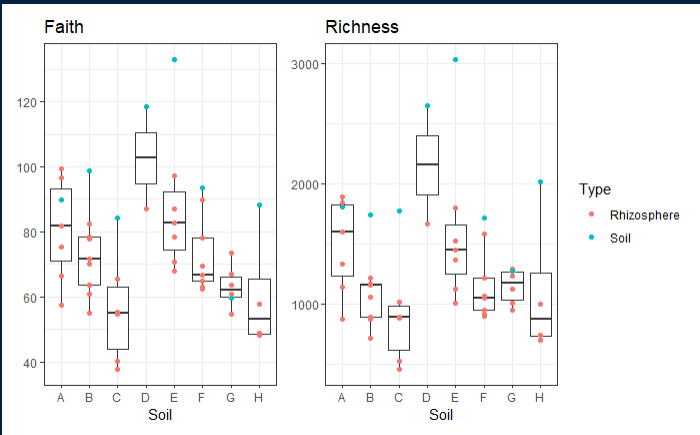

**Supplementary figure 2.** Boxplots representing the Richness (number of ASVs) and Faith’s phylogenetic diversity in the dataset according to soil of origin (A-H) and sample type (Rhizosphere or Soil).

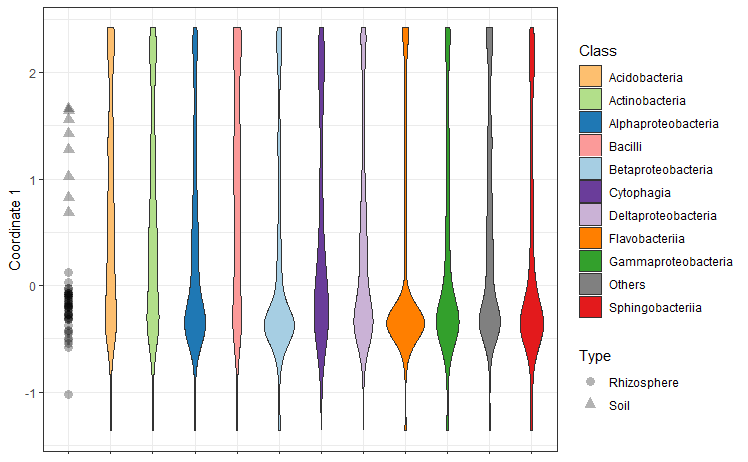
 **Supplementary figure 3.** **Not all ASVs follow the overall phylogenetic enrichment trend.** Violin plots representing the results of the PCAIV performed to explore ASV-based community composition trends after removing the patent soil of origin effect and focusing on between-type effects (permutation test, p < 0.05). Such analysis yields only one dimension, depicted in the Y-axis. The position of the samples along the axis is depicted in gray shapes, while the contributions of each ASV to the axis are merged in different density plots (violins) representing the same classes depicted in figure 2.

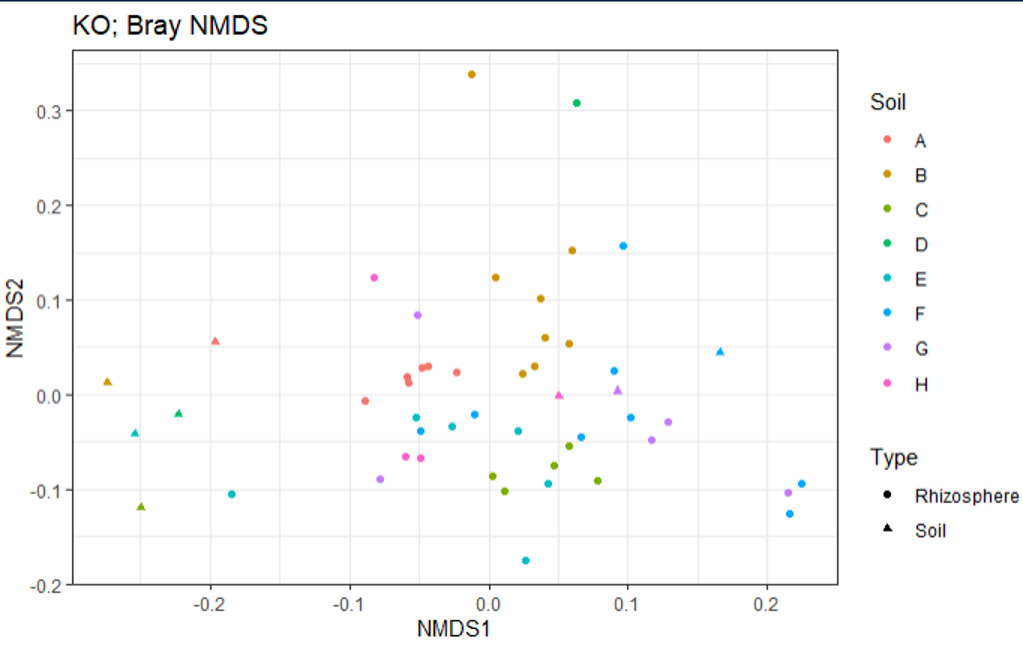

**Supplementary figure 4. Exploration of predicted functional community profiles.** The figure depicts a Non-Metric Multidimensional Scaling based on Bray-Curtis distances between predicted KO (KEGG Orthologs) compositions among samples. Samples are labeled according to soil of origin by color (A-H) and sample type by shape (Rhizosphere or Soil).

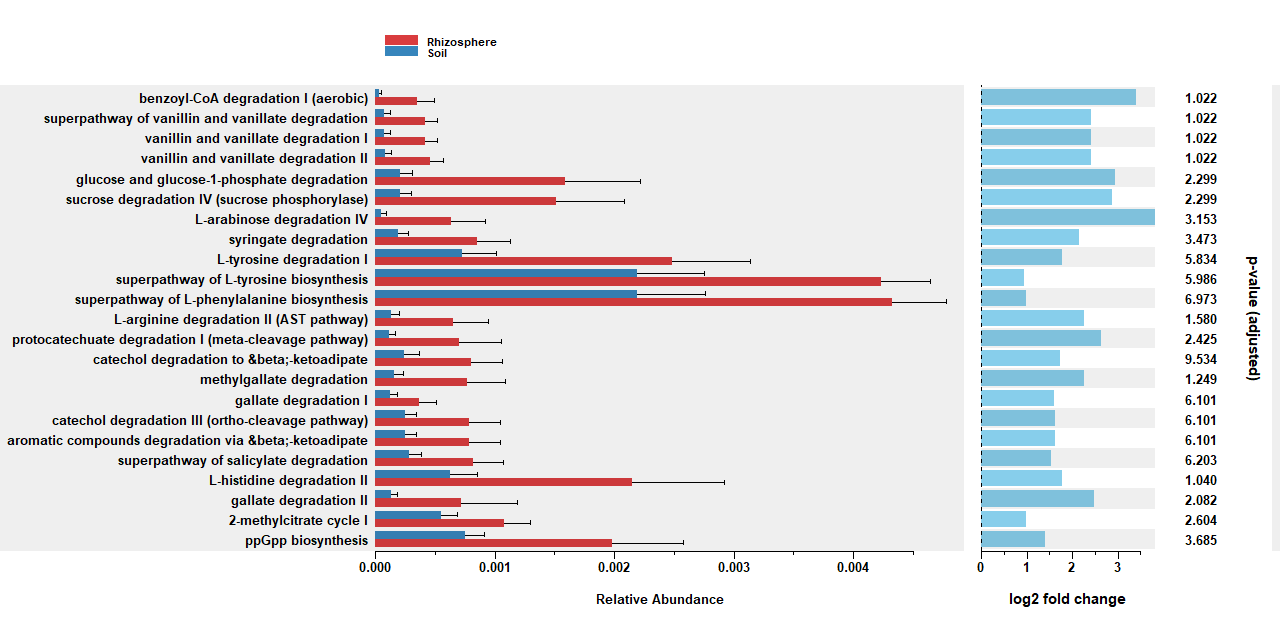

**Supplementary figure 5.** Predicted metabolic pathways enriched in rhizosphere samples (adjusted p-value<0.01).

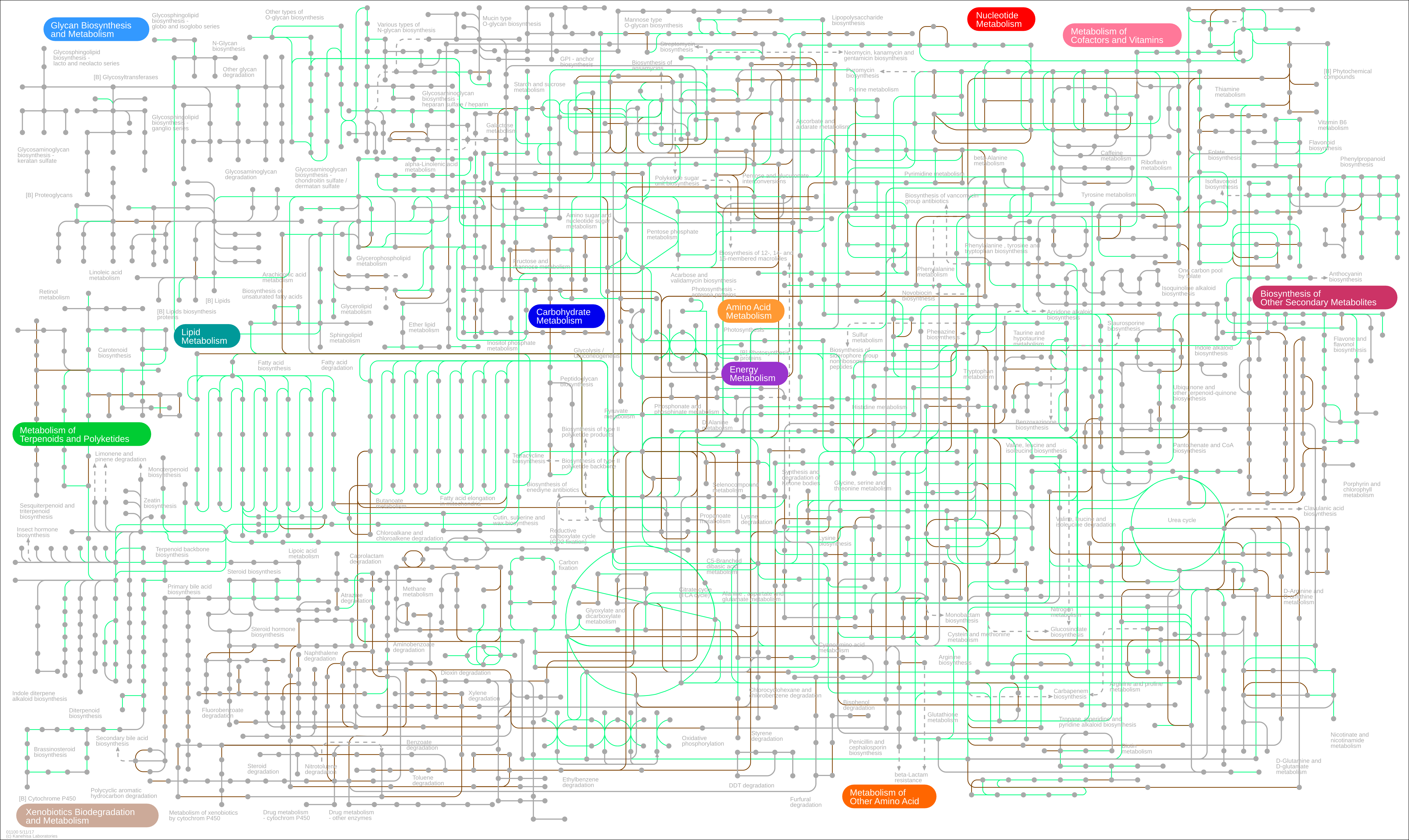

**Supplementary Figure 6. The minimal tomato rhizosphere metagenome extends its host metabolic potential.** Nodes in the map correspond to chemical compounds and edges represent enzymatic reactions. The figure provides an iPath3.0 representation of KEGG metabolic pathways, where reactions catalyzed by enzymes encoded in the tomato genome appear in green, while core reactions of the tomato rhizosphere microbiome not encoded also by the tomato genome appear in brown.

**TABLES**

| **Supplementary Table 1. Number of Kos per category exclusive for each PCG.** | | | | | | | | | | | | |
| --- | --- | --- | --- | --- | --- | --- | --- | --- | --- | --- | --- | --- |
| **Description (Pathway)** | **Node**  **119365** | **Node**  **126898** | **Node**  **147305** | **Node**  **15327** | **Node**  **19398** | **Node**  **21166** | **Node**  **28120** | **Node**  **45985** | **Node**  **52227** | **Node**  **53994** | **Node**  **55420** | **Node**  **55904** |
| Two-component system (ko02020) | 1 | 0 | 30 | 1 | 1 | 0 | 7 | 0 | 0 | 2 | 0 | 0 |
| ABC transporters (ko02010) | 0 | 0 | 9 | 3 | 4 | 0 | 7 | 0 | 0 | 0 | 0 | 11 |
| Methane metabolism (ko00680) | 0 | 1 | 2 | 1 | 0 | 0 | 1 | 0 | 0 | 11 | 0 | 0 |
| Nitrogen metabolism (ko00910) | 0 | 0 | 0 | 0 | 0 | 0 | 0 | 0 | 0 | 6 | 0 | 6 |
| Biosynthesis of siderophore group nonribosomal peptides (ko01053) | 9 | 0 | 0 | 0 | 0 | 0 | 0 | 0 | 0 | 0 | 0 | 0 |
| Vibrio cholerae pathogenic cycle (ko05111) | 0 | 0 | 0 | 0 | 0 | 5 | 1 | 0 | 0 | 0 | 0 | 0 |
| Ascorbate and aldarate metabolism (ko00053) | 0 | 1 | 1 | 0 | 1 | 1 | 0 | 0 | 1 | 0 | 0 | 0 |
| Oxidative phosphorylation (ko00190) | 0 | 0 | 5 | 0 | 0 | 0 | 0 | 0 | 0 | 0 | 0 | 0 |
| Purine metabolism (ko00230) | 0 | 0 | 3 | 0 | 0 | 1 | 1 | 0 | 0 | 0 | 0 | 0 |
| Amino sugar and nucleotide sugar metabolism (ko00520) | 0 | 0 | 1 | 0 | 0 | 1 | 1 | 0 | 0 | 0 | 0 | 2 |
| Bacterial secretion system (ko03070) | 0 | 0 | 0 | 0 | 0 | 0 | 4 | 0 | 0 | 0 | 0 | 1 |
| Cyanoamino acid metabolism (ko00460) | 0 | 0 | 0 | 0 | 0 | 0 | 3 | 0 | 0 | 1 | 0 | 0 |
| Glycerophospholipid metabolism (ko00564) | 0 | 1 | 1 | 0 | 0 | 0 | 1 | 0 | 0 | 0 | 1 | 0 |
| Cell cycle - Caulobacter (ko04112) | 0 | 0 | 0 | 0 | 0 | 0 | 0 | 0 | 0 | 4 | 0 | 0 |
| Sulfur relay system (ko04122) | 0 | 0 | 0 | 0 | 0 | 0 | 4 | 0 | 0 | 0 | 0 | 0 |
| Pentose phosphate pathway (ko00030) | 0 | 0 | 0 | 0 | 0 | 0 | 1 | 0 | 0 | 0 | 0 | 2 |
| Pentose and glucuronate interconversions (ko00040) | 0 | 0 | 1 | 0 | 0 | 0 | 0 | 0 | 1 | 0 | 1 | 0 |
| Fatty acid biosynthesis (ko00061) | 1 | 0 | 1 | 0 | 0 | 0 | 1 | 0 | 0 | 0 | 0 | 0 |
| Fatty acid metabolism (ko00071) | 0 | 1 | 1 | 0 | 0 | 0 | 1 | 0 | 0 | 0 | 0 | 0 |
| Cysteine and methionine metabolism (ko00270) | 0 | 0 | 3 | 0 | 0 | 0 | 0 | 0 | 0 | 0 | 0 | 0 |
| Lysine degradation (ko00310) | 0 | 1 | 0 | 0 | 0 | 0 | 2 | 0 | 0 | 0 | 0 | 0 |
| Phenylalanine, tyrosine and tryptophan biosynthesis (ko00400) | 0 | 0 | 2 | 0 | 0 | 0 | 0 | 0 | 1 | 0 | 0 | 0 |
| Pyruvate metabolism (ko00620) | 1 | 0 | 1 | 0 | 0 | 0 | 1 | 0 | 0 | 0 | 0 | 0 |
| Polycyclic aromatic hydrocarbon degradation (ko00624) | 0 | 0 | 0 | 2 | 0 | 0 | 0 | 0 | 0 | 0 | 0 | 1 |
| Vitamin B6 metabolism (ko00750) | 0 | 0 | 0 | 0 | 0 | 0 | 2 | 0 | 0 | 0 | 0 | 1 |
| Carotenoid biosynthesis (ko00906) | 0 | 0 | 3 | 0 | 0 | 0 | 0 | 0 | 0 | 0 | 0 | 0 |
| Glycolysis / Gluconeogenesis (ko00010) | 0 | 0 | 2 | 0 | 0 | 0 | 0 | 0 | 0 | 0 | 0 | 0 |
| Fructose and mannose metabolism (ko00051) | 0 | 0 | 1 | 0 | 0 | 0 | 0 | 0 | 0 | 0 | 0 | 1 |
| Galactose metabolism (ko00052) | 0 | 0 | 2 | 0 | 0 | 0 | 0 | 0 | 0 | 0 | 0 | 0 |
| Alanine, aspartate and glutamate metabolism (ko00250) | 1 | 0 | 0 | 0 | 0 | 0 | 0 | 0 | 0 | 0 | 1 | 0 |
| Lysine biosynthesis (ko00300) | 0 | 0 | 2 | 0 | 0 | 0 | 0 | 0 | 0 | 0 | 0 | 0 |
| Benzoate degradation (ko00362) | 0 | 1 | 0 | 1 | 0 | 0 | 0 | 0 | 0 | 0 | 0 | 0 |
| Taurine and hypotaurine metabolism (ko00430) | 0 | 0 | 0 | 0 | 0 | 0 | 0 | 0 | 0 | 0 | 0 | 2 |
| Starch and sucrose metabolism (ko00500) | 0 | 0 | 0 | 0 | 0 | 1 | 0 | 0 | 0 | 1 | 0 | 0 |
| Inositol phosphate metabolism (ko00562) | 0 | 0 | 1 | 0 | 0 | 0 | 0 | 0 | 0 | 1 | 0 | 0 |
| Glyoxylate and dicarboxylate metabolism (ko00630) | 0 | 0 | 0 | 0 | 0 | 0 | 0 | 0 | 0 | 2 | 0 | 0 |
| Porphyrin and chlorophyll metabolism (ko00860) | 0 | 0 | 1 | 0 | 0 | 0 | 0 | 0 | 0 | 1 | 0 | 0 |
| Staphylococcus aureus infection (ko05150) | 0 | 0 | 2 | 0 | 0 | 0 | 0 | 0 | 0 | 0 | 0 | 0 |
| Steroid biosynthesis (ko00100) | 0 | 0 | 1 | 0 | 0 | 0 | 0 | 0 | 0 | 0 | 0 | 0 |
| Secondary bile acid biosynthesis (ko00121) | 0 | 0 | 0 | 0 | 0 | 0 | 0 | 0 | 0 | 1 | 0 | 0 |
| Ubiquinone and other terpenoid-quinone biosynthesis (ko00130) | 0 | 0 | 1 | 0 | 0 | 0 | 0 | 0 | 0 | 0 | 0 | 0 |
| Steroid hormone biosynthesis (ko00140) | 1 | 0 | 0 | 0 | 0 | 0 | 0 | 0 | 0 | 0 | 0 | 0 |
| Glycine, serine and threonine metabolism (ko00260) | 0 | 1 | 0 | 0 | 0 | 0 | 0 | 0 | 0 | 0 | 0 | 0 |
| Geraniol degradation (ko00281) | 0 | 0 | 0 | 0 | 0 | 0 | 1 | 0 | 0 | 0 | 0 | 0 |
| Valine, leucine and isoleucine biosynthesis (ko00290) | 0 | 0 | 0 | 0 | 0 | 0 | 0 | 0 | 0 | 1 | 0 | 0 |
| beta-Lactam resistance (ko00312) | 0 | 0 | 1 | 0 | 0 | 0 | 0 | 0 | 0 | 0 | 0 | 0 |
| Arginine and proline metabolism (ko00330) | 0 | 0 | 0 | 0 | 0 | 0 | 1 | 0 | 0 | 0 | 0 | 0 |
| Tyrosine metabolism (ko00350) | 0 | 0 | 0 | 0 | 0 | 0 | 0 | 0 | 0 | 1 | 0 | 0 |
| Phenylalanine metabolism (ko00360) | 1 | 0 | 0 | 0 | 0 | 0 | 0 | 0 | 0 | 0 | 0 | 0 |
| Chlorocyclohexane and chlorobenzene degradation (ko00361) | 0 | 0 | 0 | 0 | 0 | 0 | 0 | 0 | 0 | 0 | 0 | 1 |
| Bisphenol degradation (ko00363) | 0 | 0 | 0 | 0 | 0 | 0 | 0 | 0 | 0 | 0 | 0 | 1 |
| Fluorobenzoate degradation (ko00364) | 0 | 1 | 0 | 0 | 0 | 0 | 0 | 0 | 0 | 0 | 0 | 0 |
| Tryptophan metabolism (ko00380) | 0 | 0 | 1 | 0 | 0 | 0 | 0 | 0 | 0 | 0 | 0 | 0 |
| Novobiocin biosynthesis (ko00401) | 0 | 0 | 0 | 0 | 0 | 0 | 0 | 0 | 0 | 1 | 0 | 0 |
| Phosphonate and phosphinate metabolism (ko00440) | 0 | 0 | 0 | 0 | 0 | 0 | 0 | 0 | 0 | 0 | 0 | 1 |
| Glutathione metabolism (ko00480) | 0 | 0 | 0 | 0 | 0 | 0 | 1 | 0 | 0 | 0 | 0 | 0 |
| Glycosaminoglycan degradation (ko00531) | 1 | 0 | 0 | 0 | 0 | 0 | 0 | 0 | 0 | 0 | 0 | 0 |
| Lipopolysaccharide biosynthesis (ko00540) | 0 | 0 | 0 | 0 | 0 | 0 | 1 | 0 | 0 | 0 | 0 | 0 |
| Glycerolipid metabolism (ko00561) | 0 | 1 | 0 | 0 | 0 | 0 | 0 | 0 | 0 | 0 | 0 | 0 |
| Toluene degradation (ko00623) | 0 | 0 | 0 | 0 | 0 | 0 | 0 | 0 | 1 | 0 | 0 | 0 |
| Aminobenzoate degradation (ko00627) | 0 | 0 | 0 | 0 | 0 | 0 | 0 | 0 | 0 | 1 | 0 | 0 |
| Nitrotoluene degradation (ko00633) | 0 | 0 | 0 | 0 | 0 | 1 | 0 | 0 | 0 | 0 | 0 | 0 |
| C5-Branched dibasic acid metabolism (ko00660) | 1 | 0 | 0 | 0 | 0 | 0 | 0 | 0 | 0 | 0 | 0 | 0 |
| Carbon fixation pathways in prokaryotes (ko00720) | 0 | 1 | 0 | 0 | 0 | 0 | 0 | 0 | 0 | 0 | 0 | 0 |
| Thiamine metabolism (ko00730) | 0 | 0 | 0 | 0 | 1 | 0 | 0 | 0 | 0 | 0 | 0 | 0 |
| Folate biosynthesis (ko00790) | 0 | 0 | 0 | 0 | 0 | 0 | 1 | 0 | 0 | 0 | 0 | 0 |
| Atrazine degradation (ko00791) | 0 | 1 | 0 | 0 | 0 | 0 | 0 | 0 | 0 | 0 | 0 | 0 |
| Terpenoid backbone biosynthesis (ko00900) | 0 | 0 | 1 | 0 | 0 | 0 | 0 | 0 | 0 | 0 | 0 | 0 |
| Sulfur metabolism (ko00920) | 0 | 0 | 0 | 0 | 1 | 0 | 0 | 0 | 0 | 0 | 0 | 0 |
| Isoflavonoid biosynthesis (ko00943) | 0 | 0 | 1 | 0 | 0 | 0 | 0 | 0 | 0 | 0 | 0 | 0 |
| Aminoacyl-tRNA biosynthesis (ko00970) | 0 | 1 | 0 | 0 | 0 | 0 | 0 | 0 | 0 | 0 | 0 | 0 |
| Drug metabolism - other enzymes (ko00983) | 0 | 0 | 0 | 0 | 0 | 0 | 1 | 0 | 0 | 0 | 0 | 0 |
| Biosynthesis of type II polyketide backbone (ko01056) | 1 | 0 | 0 | 0 | 0 | 0 | 0 | 0 | 0 | 0 | 0 | 0 |
| Nucleotide excision repair (ko03420) | 0 | 0 | 0 | 1 | 0 | 0 | 0 | 0 | 0 | 0 | 0 | 0 |
| Mismatch repair (ko03430) | 0 | 0 | 0 | 0 | 0 | 0 | 0 | 0 | 0 | 1 | 0 | 0 |
| Protein processing in endoplasmic reticulum (ko04141) | 0 | 0 | 0 | 0 | 0 | 0 | 0 | 0 | 0 | 1 | 0 | 0 |
| Plant-pathogen interaction (ko04626) | 0 | 0 | 0 | 0 | 0 | 0 | 1 | 0 | 0 | 0 | 0 | 0 |
| Bile secretion (ko04976) | 0 | 0 | 0 | 0 | 0 | 0 | 0 | 0 | 0 | 1 | 0 | 0 |
| African trypanosomiasis (ko05143) | 0 | 0 | 0 | 0 | 0 | 0 | 1 | 0 | 0 | 0 | 0 | 0 |
| Malaria (ko05144) | 1 | 0 | 0 | 0 | 0 | 0 | 0 | 0 | 0 | 0 | 0 | 0 |
